# Long single-molecule reads can resolve the complexity of the Influenza virus composed of rare, closely related mutant variants

**DOI:** 10.1101/036392

**Authors:** Alexander Artyomenko, Nicholas C Wu, Serghei Mangul, Eleazar Eskin, Ren Sun, Alex Zelikovsky

## Abstract

As a result of a high rate of mutations and recombination events, an RNA-virus exists as a heterogeneous “swarm” of mutant variants. The long read length offered by single-molecule sequencing technologies allows each mutant variant to be sequenced in a single pass. However, high error rate limits the ability to reconstruct heterogeneous viral population composed of rare, related mutant variants. In this paper, we present 2SNV, a method able to tolerate the high error-rate of the single-molecule protocol and reconstruct mutant variants. 2SNV uses linkage between single nucleotide variations to efficiently distinguish them from read errors. To benchmark the sensitivity of 2SNV, we performed a single-molecule sequencing experiment on a sample containing a titrated level of known viral mutant variants. Our method is able to accurately reconstruct clone with frequency of 0.2% and distinguish clones that differed in only two nucleotides distantly located on the genome. 2SNV outperforms existing methods for full-length viral mutant reconstruction. The open source implementation of 2SNV is freely available for download at http://alan.cs.gsu.edu/NGS/?q=content/2snv

## Introduction

Majority of the emerging and re-emerging diseases (influenza, hantaviruses, Ebola virus, and Nipah virus), which represent a global threat to the public health, are caused by RNA viruses [28]. RNA viruses can be featured by their robust adaptability and evolvability due to their high mutation rates and rapid replication cycles [6,15]. This enables a within-host RNA virus population to organize as a complex and dynamic mutant swarm of many highly similar viral genomes. This mutant spectrum, also known as quasispecies [9], is continuously maintained and regenerated during viral infection [7,19]. Deep sequencing has provided a new lens to monitor individual viral variants accelerating the understanding of escape and resistance mechanisms [3,26], in addition to providing insights about the viral evolutionary landscape and the genomic interactions [22,30,39].

Short reads offered by commonly used fragmentation-based protocols protocols are well suited to detect discrete genome components, such as the frequency of each single-nucleotide polymorphism. However, high similarity of the individual viral genomes imposes a huge challenge to assemble discrete components into a population of full-length viral genomes. In particular, mutations are often located on the distances unreachable by the short reads. Therefore even hybrid technologies based on error correction of PacBio reads with Illumina reads were not applied to sequencing of viral variants. Indeed, short reads cannot tell the allele - the same short read is equally well mapped to a variant with the major allele and a variant with the minor allele.

Single Molecule Real Time (SMRT) sequencing is a parallelized single molecule DNA sequencing method. PacBio SMRT sequencing reads are much longer than sequencing reads provided by Illumina, however, its throughput is much lower and the error rate is significantly higher. The read length offered by a singlemolecule sequencing protocol [8] is comparable to the genome size of most RNA viruses. It allows each genome variant to be sequenced in a single pass, providing an accurate phasing of the distant mutations. The main drawbacks of the long single-molecule technologies are the high error rate and comparatively low throughput, limiting ability of those technologies to study the heterogeneous viral populations. Thus, a complete profiling of all viral genomes within a mutant spectrum is not yet possible.

Recently, this problem has been addressed using various computational and statistical approaches implemented in Quasirecomb [36], PredictHaplo [32], HaploClique [35], VGA [24], and *k*GEM [33]. These methods perform reasonably well on short reads with high coverage and low error rate, but our experimental validation shows far from satisfactory performance on the sequencing data provided by single-molecule technologies. Also a workflow for reconstruction of closely related variants from raw reads generated during SMRT sequencing was proposed in [4]. Note that a recent method for haplotyping using Pacbio reads proposed in [34] is only applicable for diploid organisms and is not suitable for viral haplotyping with numerous variants.

In this paper, we present two Single Nucleotide Variants (2SNV), a comprehensive method for the accurate reconstruction of the heterogeneous viral population from the long single-molecule reads. The 2SNV method hierarchically clusters together reads containing pairs of correlated (i.e., linked) SNVs until no cluster has correlated SNVs left and outputs consensus of each cluster. It allows to reduce error rate and differentiate true biological variants from sequencing artifacts, thus providing increased accuracy to study diversity and composition of the viral spectrum. To benchmark the sensitivity of 2SNV, we performed a single-molecule sequencing experiment on a sample containing a titrated level of known viral mutant variants. We were able to reconstruct a haplotype with a frequency of 0.2% and distinguish clones that differed in only two nucleotides. We also showed that 2SNV outperformed existing haplotype reconstruction tools. With a high sensitivity and accuracy, 2SNV is anticipated to facilitate not only viral quasispecies reconstruction, but also other biological questions that require detection of rare haplotypes such as genetic diversity in cancer cell population, and monitoring B-cell and T-cell receptor repertoire.

## Methods

Any method for reconstruction of viral variants from single-molecule reads should overcome low volume and high error rate of sequencing data combined with very high similarity and very low frequency of viral variants. This challenge is equivalent to extraction of an extremely weak signal from very noisy background with signal-to-noise ratio approaching zero. However impossible this task may seem, a satisfactory solution can be based on distinguishing randomness of the noise from systematic signal repetition. Previously, linkage between SNVs was used for distinguishing sequencing errors from SNVs [23], however, to the best of our knowledge, it was never applied for haplotyping.

Since all reads are from the same RNA region of very similar sequences, they can be reliably aligned to each other. In general, the errors in different positions are independent from each other and the further these positions are from each other the less likely any dependency can be caused by systematic errors. Therefore, even slightly more than expected co-occurrence of two rare alleles in non-adjacent positions may serve as a trustful signature of one or more rare variants having the both rare alleles. Such single nucleotide variations (SNVs) are called linked.

The proposed 2SNV method recursively clusters reads containing pairs of linked SNVs until no pair of SNVs exhibits statistically significant linkage in any cluster. Then each cluster should contain just a single viral variant which can be simply reconstructed as the consensus of all reads in the cluster.

In the remainder of the section we derive statistical conditions of SNV linkage and then give detailed description of the 2SNV method which identifies rare variants based SNV pairs satisfying these conditions.

### Linkage of SNV pairs

In this section we analyze statistical significance of the linkage between a pair of SNVs which allows to distinguish reads emitted by a rare variant from background errors.

We assume that errors are random and a rare variant has at least 2 mismatches with other variants. Let us consider an arbitrary pair of two distinct positions *I*, *J* ∈ {1,…, *L*}, *I* ≠ *J*, where *L* be the length of the amplicon (see Figure 1b). Let *I*_1_ and *J*_1_ be the alleles of the most frequent 2-haplotype (*I*_1_ *J*_1_). Note that (*I*_1_ *J*_1_) should be a 2-haplotype from at least one true viral variant assuming that the error rates in the *I*-th and *J*-th positions are small and independent.

**Fig. 1.**
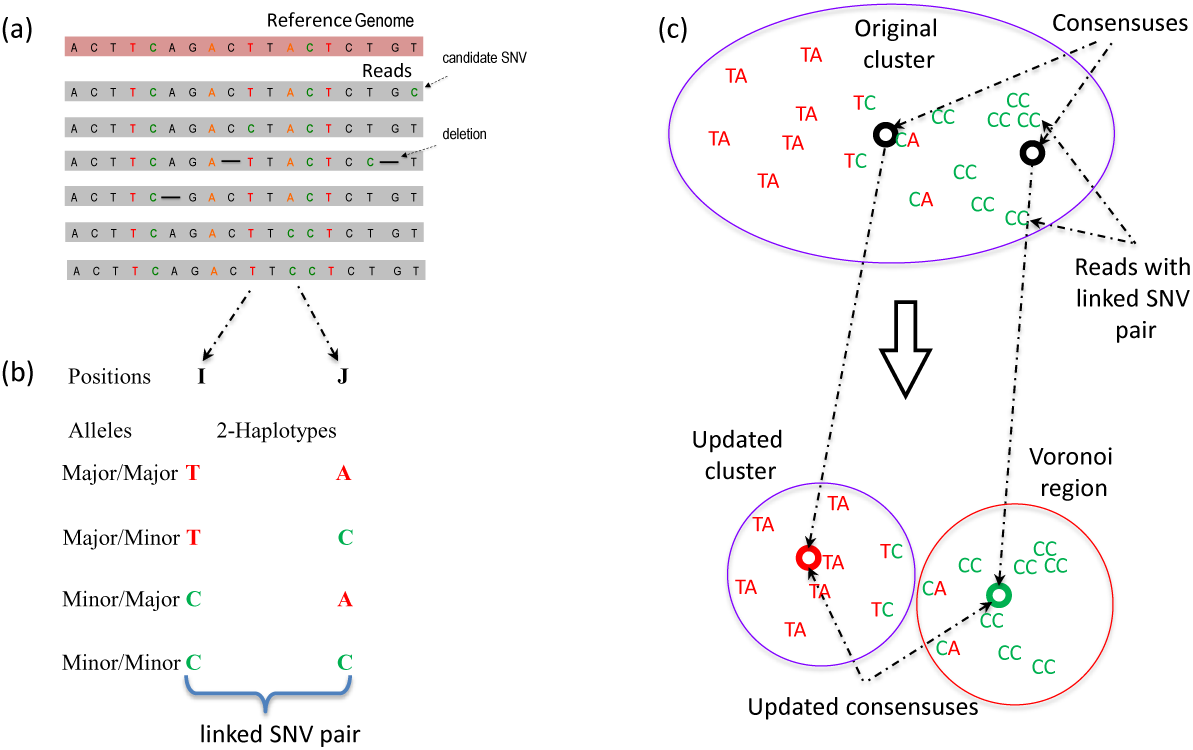
Overview of the 2SNV method: (a) Multiple sequence alignment of reads from the same amplicon; (b) Identification of a linked SNV pair in positions *I* and *J*; (c) Recursive cluster splitting: (i) finding consensus of reads with the linked SNV pair, (ii) finding Voronoi region of this consensus, (iii) update the original cluster and the consensuses for the two new clusters.

Let *I*_2_ ≠ *I*_1_ and *J*_2_ ≠ *J*_1_ be the alleles of another 2-haplotype. Let *E*_*kl*_, *k*, *l* ∈ {1, 2}, be the expected number of reads with 2-haplotypes (*I*_*k*_ *J*_*l*_). The following theorem can be used to decide if the haplotype *I*_2_ ≠ *I*_1_ exists.

#### Theorem 1.

Assume that the sequencing error is random, independent and does not exceed 50%. If no viral variant with the haplotype (*I*_2_ *J*_2_) exists, then the expected value of *E*_22_ is at most

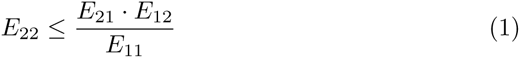

The inequality (1) becomes an equality if at least one of 2-haplotypes (*I*_1_ *J*_2_) or (*I*_2_ *J*_1_) also does not exist.

#### Proof.

Let 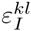 and 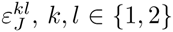, be the probabilities to observe the allele *l* instead of the true allele *k* in the positions *I* and *J*, respectively. We are not going to estimate the parameters 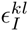. The model only assumes that these parameters are random, independent, and do not exceed 50%.

Let *T*_*kl*_, *k*, *l* ∈ {1, 2}, be the true count of 2-haplotypes (*I*_*k*_ *J*_*l*_). Then error randomness and independence imply that

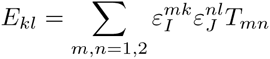

In order to prove (1), it is sufficient to show that *E*_11_ · *E*_22_ ≤ *E*_12_ · *E*_21_ assuming that *T*_22_ = 0. Indeed,

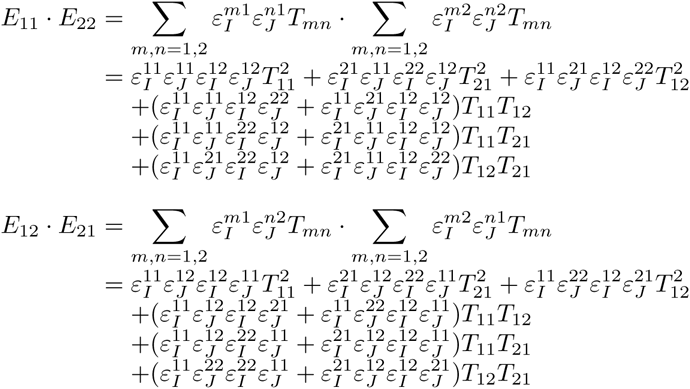

Note that only coefficients for *T*_12_*T*_21_ are different for these products. Therefore, if either *T*_12_ = 0 or *T*_21_ = 0, then *E*_11_ · *E*_22_ = *E*_12_ · *E*_21_. Otherwise, let all three 2-haplotypes (*I*_1_ *J*_1_), (*I*_1_ *J*_2_), and (*I*_2_ *J*_1_) exist. Then

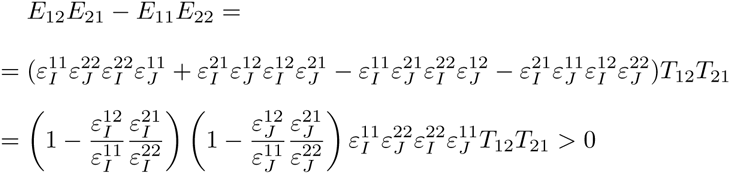

The last inequality holds since observing the true allele is more probable than observing the erroneous allele and, therefore, 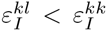 and 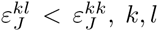 ∈ {1, 2}. QED

Note that Theorem 1 does not require linkage disequilibrium of haplotypes - the lack of linkage is explained by errors. The 2SNV method uses Theorem 1 to decide if the alleles *I*_2_ and *J*_2_ are linked as follows. Let *O*_*kl*_, *k*, *l* ∈ {1, 2}, be the observed number of reads with 2-haplotypes (*I*_*k*_ *J*_*l*_). Let *n* be the total number of *O*_22_ reads covering the both positions *I* and *J*, then

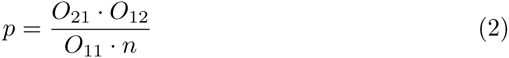

is the largest probability of observing the 2-haplotype (*I*_2_ *J*_2_) among these *n* reads. The probability to observe at least *O*_22_ reads in the (*n*, *p*) binomial distribution equals

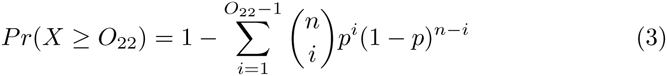

Since we are looking for a pair of SNVs among 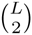 possible pairs, we also adjust to multiple testing using Bonferroni correction requiring

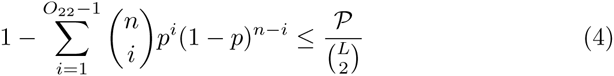

where *p* is defined in (2) and ***P*** is the user-defined *P*-value, by default ***P*** = 0.01.

Finally, when the cluster is too small, the statistical test (4) may be not stringent enough to weed out spurious linkages. Therefore, we require the number of reads *O*_22_ to be at least an empirically defined value (by default equal 30), in order to decide whether there is an additional haplotype producing these reads.

Note that the binomial model used in (4) may not be stringent enough to compensate for reducing PPV caused by overdispersion especially for higher coverage. In future releases of our tool we plan to take in account additional variance modeling unknown experimental data processes contributing to variance, e.g., replacing the binomial distribution with the beta-binomial distribution.

### 2NV method for viral variant reconstruction

The input to 2SNV consists of a set of aligned PacBio reads (see Figure 1(a)). Alignment required to be in a form of multiple sequence alignment (MSA). The MSA algorithms are too slow to handle PacBio datasets, so instead, we use pairwise alignment by BWA [20] and b2w from Shorah [41] to transform pairwise alignment to MSA format.

The main novel step of the 2SNV algorithm identifies a pair of linked SNVs (see Figure 1(b)). with higher than expected portion of reads containing the 2-haplotype with the both minor alleles according to (2–4).

The 2SNV method maintains a partition of all reads into clusters. Each cluster is assumed to consist of the reads emitted by the single variant coinciding with the cluster consensus (see Figure 1(c)). Until no pair of SNVs in the cluster *C* is linked, we recursively partition *C* into two clusters *C*_1_ and *C*_2_. *C*_1_ consists of reads with the linked pair of SNVs *C*_2_ consists of the remaining reads of *C*. We further modify *C*_1_ and *C*_2_ by replacing them with the Voronoi regions of their consensuses, where *the Voronoi region* of the consensus *c*_1_ of *C*_1_ consists of reads that are closer to *c*_1_ than to the consensus of *C*_2_. Finally, *k*GEM finds maximum likelihood estimates of frequencies of haplotypes represented by cluster consensuses using expectation-maximization algorithm [33].

Algorithm 1 describes the formal pseudocode of the 2SNV algorithm.

#### Algorithm 1 2SNV Algorithm

**procedure 1: constructing the consensus haplotype for all reads**:

Initialize the set of all clusters with a single cluster with all reads *C* ← {*R*}
For each position *i* find allele of highest frequency *a*_*i*_
*Consensus*(*C*) ← (*a*_*i*_,…, *a*_*L*_)
**procedure 2: partitioning reads into simple clusters**

**while** not all clusters are simple do
**for** each non-simple cluster *C* ∈ *C* **do**
**if** no pair SNVs is linked according to (2–4) then Regard *C* as a simple cluster
**else**
Find a pair of linked SNVs *I*_2_ and *J*_2_ minimizing (3)
Find the set *C*_1_ of all reads with the 2-haplotype (*I*_2_*J*_2_)
Find the consensus *c*_1_ ← *Consensus*(*C*_1_)
*C*_1_ ← *Voronoi*(*c*^1^)
*C*_2_ ← *C*\*C*_1_, *C*_2_ ← *Consensus*(*C*_2_)
*C*←*C*∪{*C*_1_}∪{*C*_2_}\{*C*}
**procedure 3: estimating frequencies of the consensuses of simple clusters** Run *k*GEM algorithm for the set of haplotypes {*Consensus*(*C*), *C* ∈ *C*.

## Results

We were using 3 datasets: PacBio reads from a single IAV clone and 10 IAV clones, and simulated PacBio reads from 20 HCV clones.

Error-prone PCR was performed on the influenza A virus (A/WSN/33) PB2 segment using GeneMorph II Random Mutagenesis Kits (Agilent Technologies, Westlake Village, CA) according to manufacturer’s instruction. The 2kb region was amplified from the IAV viral population and subjected to PacBio RS II sequencing using 2 SMRT cells with P4-C2. The average read length was 1973bp and ranges from 200bp to 5kb. Some reads are much longer than the amplified region due to long insertions which are sequencing errors. Raw sequencing data have been submitted to the NIH Short Read Archive (SRA) under accession number: BioProject PRJNA284802. The nucleotide sequences of the 10 clones are freely available at http://alan.cs.gsu.edu/NGS/?q=content/2snv.

**The dataset with a single IAV clone**. There total number of reads were 11,907 and the average Hamming distance between the true haplotype and reads is 14.4%.

**The dataset with 10 IAV clones**. 10 independent clones, ranging from 1 to 13 mutations from the original single were selected. These 10 clones were mixed at a geometric ratio with two-fold difference in occurrence frequency for consecutive clones starting with the maximum frequency of 50% and the minimum frequency of 0.1%. The pairwise edit distance between clones are given in the heat-map on Figure 2 in Supplement. In total, there were 33,558 reads generated from 10 clones.

**The simulated dataset with 20 HCV clones**. 21K simulated PacBio reads were generated from 1739-bp long fragment from the E1E2 region of 20 HCV sequences [38] using simulator pbsim [29]. The reads were simulated with mean accuracy 98% and minimum accuracy 95% reflecting advancements in PacBio technology. We have generated reads 10 times for two distributions of the clone frequencies – uniform (all frequencies are 5%) and skewed (a single clone has 90.5% and every other clone has frequency 0.5%).

### Reconstruction of viral variants

2SNV was compared with 2 tools originally tuned to handle HIV variants (PredictHaplo [32] and Quasirecomb [36]) and *k*GEM [33] tuned for a short HCV amplicon. We could not compare with HaploClique [35] since it is no longer maintained by the authors. A workflow [4] is not currently available and we were not able to run it on our data. Also the experimental data in [4] are also not fully available and we were not able to run 2SNV on these data.

For the dataset with a single IAV clone 2SNV, *k*GEM, and PredictHaplo were able to reconstruct no more a single variant which perfectly matches the original clone. Quasirecomb reported multiple variants none fully matching the original clone.

For the dataset with 10 IAV clones, 2SNV reported 10 haplotypes: the 9 most frequent haplotypes exactly matching 9 most frequent clones and the least frequent haplotype (1%) not matching any clone. The correlation between the estimated and true frequencies of the 9 correctly reconstructed haplotypes is 99.4%. PredictHaplo was able to reconstruct only 6 true variants missing 4 variants with total frequency of 8% while not having any false positives. In order to reliably compare the reconstruction rate of two methods, we have applied them to 40 sub-samples of the original data (each subsample consists of 33558 reads randomly selected with repetition from the original data). The results are presented on Figure 4 and Table 1 in Supplement. *k*GEM was able to reconstruct only 2 most frequent clones and Quasirecomb failed to reconstruct even a single clone.

In order to estimate how accuracy of reconstruction methods depends on the coverage, we have randomly sub-sampled *N* reads (*N* = 500, 1000, 2000, 4000, 8000, 16000) from the original 33558 reads and run 2SNV and PredictHaplo. The results are shown on Figure 2 in Supplement. For each coverage and each clone (except Clone5), 2SNV more accurately estimates the frequency. Clone6 and Clone8 for all sub-samples, Clone4 for *N* ≤ 8000 and Clone 3 for *N* ≤ 1000 are missed by PredictHaplo but reconstructed by 2SNV. Clone6 which is only two mutations away from the more frequent Clone5 was successfully reconstructed for *N* ≥ 4000 while PredictHaplo was never able to reconstruct Clone6. Note that since these 2 SNVs between Clone5 and Clone6 are far apart, only long reads can reconstruct this rare variant. From the last plot one can see that the false positive rate for PredictHaplo is also higher than for 2SNV, e.g. 2SNV does not report false positives for *N* ≤ 8000. The averages of all runs are given in Table 2 in Supplement.

For the simulated dataset with 20 HCV variants, we have compared 2SNV only with PredictHaplo. For the uniform frequency distribution the average sensitivity and PPV for 2SNV are 85% and 100%, respectively, while for PredictHaplo the corresponding values are 72% and 53%, respectively. For the skewed frequency distribution, the average sensitivity and PPV for 2SNV are 99% and 69%, respectively, while for PredictHaplo the corresponding values are 36% and 46%, respectively.

**Runtime**. The runtime of 2SNV is linear with respect to the number of reads, however implementation is *O(nlogn)* due to parallelization (see Figure 5 in Supplement) and quadratic with respect to the length of the amplicon region. For all experiments we used the same PC (Intel(R) Xeon(R) CPU X5550 2.67GHz x2 8 cores per CPU, DIMM DDR3 1333 MHz RAM 4Gb x12) with operating system CentOS 6.4.

## Discussion

Haplotype phasing represents one of the biggest challenges in next-generation sequencing due to the short read length. The recent development of single-molecule sequencing platform produces reads that are sufficiently long to span the entire gene or small viral genome. It not only benefits the assembly of genomic regions with tandem repeat [5,18,37], but also offers the opportunity to examine the genetic linkage between mutations. In fact, it is shown that the long read in single-molecule sequencing aids haplotype phasing in diploid genome [31], and in polyploid genome [1]. Nonetheless, the sequencing error rate of singlemolecule sequencing platform is extremely high (≈ 14% as estimated by this study), which hampers its ability to reconstruct rare haplotypes. This drawback prohibits single-molecule sequencing platform from applications in which a high sensitivity of haplotypes are needed, such as quasispecies reconstruction. In this study, we have developed 2SNV, which allows quasispecies reconstruction using single-molecule sequencing despite the high sequencing error rate. The high sensitivity of 2SNV permits the detection of extremely rare haplotypes and distinguish between closely related haplotypes. Based on titrated levels of known haplotypes, we demonstrates that 2SNV is able to detect a haplotype that has a frequency as low as 0.2%. This sensitivity is comparable to many deep sequencing-based point mutation detection methods [10,11,13,21]. In addition, 2SNV successfully distinguishes between Clone5 and Clone6 in this study, which are only two nucleotides away from each other. It highlights the sensitivity of 2SNV to distinguish closely related haplotypes. Our results also show that the sensitivity is coverage-dependent, implying that the sensitivity of 2SNV may further improve when sequencing depth increases. Therefore, the constant increase of sequencing throughput offered by single-molecule sequencing technology provides the unprecedented resolution promising to increase number of discovered rare haplotypes.

The ability to accurately determine the genomic composition of the viral populations and identify closely related viral genomes makes our tool applicable for dissecting evolutionary trajectories and examining mutation interactions in RNA viruses. Evolutionary trajectories and mutation interactions have been shown to play an important role in viral evolution, such as drug resistance [2,3,26,39], immune escape [12], and cross-species adaptation [14,16]. An unbiased and accurate understanding of the genomic composition of the RNA viruses opens a new avenue to study the underlying mechanism of adaptation, persistence and virulence factors of the pathogen, which are yet to be comprehended.

While viral quasispecies reconstruction is used as a proof-of-concept in this study, the application of 2SNV can be extended to detect haplotype variants in any sample with high genetic heterogeneity and diversity, such as B-cell and T-cell receptor repertoire, cancer cell populations, and metagenomes. It is shown that monitoring B-cell and T-cell receptor repertoire helps investigate virus-host interaction dynamics [17,27,40,42,43]. Furthermore, examining the genetic composition of the cancer cell populations in high sensitivity can facilitate diagnosis and treatment [25]. Therefore, we anticipate that 2SNV will benefit different subfields of biomedical research in the genomic era. We also propose that 2SNV can be applied to increase the resolution of metagenomics profiling from species level to strain level. In summary, 2SNV is a widely applicable tool as single-molecule sequencing technology being popularized.

## Supplement.

The Supplement to this paper containing Figures 1-6 and Tables 1-2 is available here: http://alan.cs.gsu.edu/NGS/?q=content/2snv_supplement

## Acknowledgments.

We would like to thank H. Hao for performing the PacBio sequencing at Johns Hopkins Deep Sequencing & Microarray Core Facility. A.A. was supported by GSU Molecular Basis of Disease Fellowship. S.M. and E.E were supported by National Science Foundation grants 0513612, 0731455, 0729049, 0916676, 1065276, 1302448 and 1320589, and National Institutes of Health grants K25-HL080079, U01-DA024417, P01-HL30568, P01-HL28481, R01-GM083198, R01-MH101782 and R01-ES022282. S.M. was supported in part by Institute for Quantitative & Computational Biosciences Fellowship, UCLA.

## References

1. Aguiar, D. Istrail, S.: Haplotype assembly in polyploid genomes and identical by descent shared tracts. Bioinformatics 29(13), i352–360 (Jul 2013)

2. Beerenwinkel, N. et al.: Diversity and complexity of hiv-1 drug resistance: a bioinformatics approach to predicting phenotype from genotype. Proceedings of the National Academy of Sciences 99(12), 8271–8276 (2002)

3. Bushman, F.D. et al.: Massively parallel pyrosequencing in hiv research. Aids 22(12), 1411–1415 (2008)

4. Dilernia, D.A. et al.: Multiplexed highly-accurate DNA sequencing of closely-related HIV-1 variants using continuous long reads from single molecule, real-time sequencing. Nucleic Acids Research (2015)

5. Doi, K. et al.: Rapid detection of expanded short tandem repeats in personal genomics using hybrid sequencing. Bioinformatics 30(6), 815–822 (Mar 2014)

6. Domingo, E.: Mutation rates and rapid evolution of RNA viruses. The evolutionary biology of viruses pp. 161–184 (1994)

7. Domingo, E. Holland, J.: RNA virus mutations and fitness for survival. Annual Reviews in Microbiology 51(1), 151–178 (1997)

8. Eid, J. et al.: Real-time dna sequencing from single polymerase molecules. Science 323(5910), 133–138 (2009)

9. Eigen, M.: Selforganization of matter and the evolution of biological macromolecules. Naturwissenschaften 58(10), 465–523 (1971)

10. Flaherty, P. et al.: Ultrasensitive detection of rare mutations using next-generation targeted resequencing. Nucleic Acids Res. 40(1), e2 (Jan 2012)

11. Forshew, T. et al.: Noninvasive identification and monitoring of cancer mutations by targeted deep sequencing of plasma DNA. Sci Transl Med 4(136), 136ra68 (May 2012)

12. Goepfert, P.A. et al.: Transmission of hiv-1 gag immune escape mutations is associated with reduced viral load in linked recipients. The Journal of experimental medicine 205(5), 1009–1017 (2008)

13. Harismendy, O. et al.: Detection of low prevalence somatic mutations in solid tumors with ultra-deep targeted sequencing. Genome Biol. 12(12), R124 (2011)

14. Herfst, S. et al.: Airborne transmission of influenza a/h5n1 virus between ferrets. Science 336(6088), 1534–1541 (2012)

15. Holland, J. et al.: Rapid evolution of RNA genomes. Science 215(4540), 1577–1585 (1982)

16. Imai, M. et al.: Experimental adaptation of an influenza h5 ha confers respiratory droplet transmission to a reassortant h5 ha/h1n1 virus in ferrets. Nature 486(7403), 420–428 (2012)

17. Klarenbeek, P.L. et al.: Deep sequencing of antiviral T-cell responses to HCMV and EBV in humans reveals a stable repertoire that is maintained for many years. PLoS Pathog. 8(9), e1002889 (Sep 2012)

18. Schrago, C.G. Carvalho, A.B.: Long-Read Single Molecule Sequencing to Resolve Tandem Gene Copies: The Mst77Y Region on the Drosophila melanogaster Y Chromosome. G3 (Bethesda) 5(6), 1145–1150 (2015)

19. Lauring, A.S. Andino, R.: Quasispecies theory and the behavior of rna viruses. PLoS Pathogens 6(7), e1001005 (2010)

20. Li, H. Durbin, R.: Fast and accurate short read alignment with burrows-wheeler transform. Bioinformatics 25(14), 1754–1760 (2009)

21. Li, M. Stoneking, M.: A new approach for detecting low-level mutations in next-generation sequence data. Genome Biol. 13(5), R34 (2012)

22. Liu, J. et al.: Analysis of low-frequency mutations associated with drug resistance to raltegravir before antiretroviral treatment. Antimicrobial agents and chemotherapy 55(3), 1114–1119 (2011)

23. Macalalad, A.R. et al.: Highly sensitive and specific detection of rare variants in mixed viral populations from massively parallel sequence data. PLoS Comput Biol 8(3), e1002417 (2012)

24. Mangul, S. et al.: Accurate viral population assembly from ultra-deep sequencing data. Bioinformatics 30(12), i329–i337 (2014)

25. Mardis, E.R. Wilson, R.K.: Cancer genome sequencing: a review. Hum. Mol. Genet. 18(R2), R163–168 (Oct 2009)

26. Margeridon-Thermet, S. et al.: Ultra-deep pyrosequencing of hepatitis b virus quasispecies from nucleoside and nucleotide reverse-transcriptase inhibitor (nrti)-treated patients and nrti-naive patients. Journal of Infectious Diseases 199(9), 1275–1285 (2009)

27. Miconnet, I.: Probing the T-cell receptor repertoire with deep sequencing. Curr Opin HIV AIDS 7(1), 64–70 (Jan 2012)

28. Murphy, F.A. Kingsbury, D.W.: Virus taxonomy. Fields Virology 2, 15–57 (1996)

29. Asai, K. Hamada, M.: Pbsim: Pacbio reads simulator toward accurate genome assembly. Bioinformatics 29(1), 119–121 (2013)

30. Palmer, S. et al.: Selection and persistence of non-nucleoside reverse transcriptase inhibitor-resistant HIV-1 in patients starting and stopping non-nucleoside therapy. Aids 20(5), 701–710 (2006)

31. Pendleton, M. et al.: Assembly and diploid architecture of an individual human genome via single-molecule technologies. Nat. Methods (Jun 2015)

32. Beerenwinkel, N. Roth, V.: HIV haplotype inference using a propagating Dirichlet process mixture model. IEEE/ACM Transactions on Computational Biology and Bioinformatics (TCBB) 11(1), 182–191 (2014)

33. Skums, P. et al.: Computational framework for next-generation sequencing of heterogeneous viral populations using combinatorial pooling. Bioinformatics 31(5), 682–690 (2015)

34. Sharon, D. Snyder, M.P.: Defining a personal, allele-specific, and single-molecule long-read transcriptome. Proceedings of the National Academy of Sciences 111(27), 9869–9874 (2014)

35. Töpfer, A. et al.: Viral quasispecies assembly via maximal clique enumeration. In: RECOMB. pp. 309–310 (2014)

36. Töpfer, A. et al.: Probabilistic inference of viral quasispecies subject to recombination. Journal of Computational Biology 20(2), 113–123 (2013)

37. Ummat, A. Bashir, A.: Resolving complex tandem repeats with long reads. Bioinformatics 30(24), 3491–3498 (Dec 2014)

38. Von Hahn, T. et al.: Hepatitis c virus continuously escapes from neutralizing antibody and t-cell responses during chronic infection in vivo. Gastroenterology 132(2), 667–678 (2007)

39. Ronaghi, M. Shafer, R.: Characterization of mutation spectra with ultra-deep pyrosequencing: Application to HIV-1 drug resistance. Genome Research 17(8), 1195–1201 (2007)

40. Wu, X. et al.: Focused evolution of HIV-1 neutralizing antibodies revealed by structures and deep sequencing. Science 333(6049), 1593–1602 (Sep 2011)

41. Eriksson, N. Beerenwinkel, N.: Shorah: estimating the genetic diversity of a mixed sample from next-generation sequencing data. BMC Bioinformatics 12(1), 119 (2011)

42. Zhu, J. et al.: Mining the antibodyome for HIV-1-neutralizing antibodies with next-generation sequencing and phylogenetic pairing of heavy/light chains. Proc. Natl. Acad. Sci. U.S.A. 110(16), 6470–6475 (Apr 2013)

43. Zhu, J. et al.: De novo identification of VRC01 class HIV-1-neutralizing antibodies by next-generation sequencing of B-cell transcripts. Proc. Natl. Acad. Sci. U.S.A. 110(43), E4088–4097 (Oct 2013)

